# The Interplay of Furin Cleavage and D614G in Modulating SARS-CoV-2 Spike Protein Dynamics

**DOI:** 10.1101/2025.01.27.635166

**Authors:** Sophie R. Shoemaker, Megan Luo, Kim-Marie A. Dam, John E. Pak, Magnus A. G. Hoffmann, Susan Marqusee

## Abstract

We report a detailed analysis of the full-length SARS-CoV-2 spike dynamics within a native-like membrane environment and variants inaccessible to studies on soluble constructs by conducting hydrogen-deuterium exchange mass spectrometry (HDX-MS) on enveloped virus-like particles (eVLPs) displaying various spike constructs. We find that the previously identified open-interface trimer conformation is sampled in all eVLP-displayed spike variants studied including sequences from engineered vaccine constructs and native viral sequences. The D614G mutation, which arose early in the pandemic, favors the canonical ‘closed-interface’ prefusion conformation, potentially mitigating premature S1 shedding in the presence of a cleaved furin site and providing an evolutionary advantage to the virus. Remarkably, furin cleavage at the S1/S2 boundary allosterically increases the flexibility of the S2’ site, which may facilitate increased TMPRSS2 processing, enhancing viral infectivity. The use of eVLPs in HDX-MS studies provides a powerful platform for studying viral and membrane proteins in near-native environments.

## Introduction

SARS-CoV-2 has profoundly affected society since late 2019. This betacoronavirus infects host cells through the spike protein, which decorates the surface of the enveloped virus. The spike protein is responsible for recognizing the host cell receptor, angiotensin-converting enzyme 2 (ACE2)^1,2^. Following cleavage at the S2’ protease site by TMPRSS2^1^, this viral-fusion protein undergoes dramatic conformational changes, ultimately leading to membrane fusion of the host and viral membranes. As the largest surface protein on the virus with a crucial role in the infection process, the spike protein is the primary target for vaccines and therapeutics^3,4^.

The evolution of SARS-CoV-2 has been marked by several sequence changes in the spike protein that have significantly impacted transmissibility and virulence. The acquisition of a multibasic furin cleavage site at the S1/S2 junction (R-R-A-R), a distinctive characteristic of SARS-CoV-2 compared to other closely related betacoronaviruses, has been shown to be required for human lung infection^5^. Proteolytic activation of spike at the S1/S2 junction is thought to play a role in viral pathogenesis and tropism^6^. Other related coronaviruses have different protease cleavage sites which are processed with variable efficiency. For instance, Middle-East respiratory syndrome (MERS) coronavirus has a minimal furin cleavage site (R-X-X-R)^7^, and SARS-CoV-1 has a single arginine at this junction^5,8^. Notably, the SARS-CoV-2 spike furin cleavage site has an additional basic residue, making it more similar to the optimal furin cleavage site (R-R-A-R *vs* R-X-R-R), and has been shown to be cleaved more efficiently than the S1/S2 junction of SARS-CoV-1 and MERS^8^. It is proposed that efficient cleavage at the S2’ protease site by TMPRSS2 on the host cell surface—triggering fusion— requires SARS-CoV-2 spike protein to first be pre-processed by furin at the S1/S2 junction^5,8^. To date, no molecular mechanism has been identified to explain how cleavage at the S1/S2 protease site enhances processing by TMPRSS2 at the S2’ protease site. The majority of soluble spike constructs developed for biochemical and structural studies as well as some vaccine constructs^4^, have an ablated furin cleavage site. The impact of this deletion and the resulting lack of any potential cleavage at this site on the structure and antigenicity of these constructs remains unknown.

Another notable sequence feature of the SARS-CoV-2 spike is the early emergence and fixation of the D614G mutation, which resides in the S1 domain of the spike protein. In January 2020, viral genomes revealed a high frequency of the A23403G single nucleotide polymorphism, resulting in the change of an aspartic acid to a glycine at position 614 (D614G) of the spike protein^9,10^. By June 2020, this mutation had reached near fixation globally^9,10^ and has persisted in all subsequent variants. Studies in various cell types, using both authentic SARS-CoV-2 and viral pseudotypes, have shown that the G614 variant is 2- to 9-fold more infectious than D614^10–12^. The molecular mechanism of the evolutionary advantage for the D614G mutation is unclear. Experiments have shown that it is not due to immune evasion, spike expression and processing, nor viral assembly^10–13^. Several competing models have been proposed for why D614G provides a competitive advantage for the virus including reducing premature S1 dissociation from S2^12,14^ and increasing the proportion of RBDs in the up conformation^10^. Given conflicting data, it remains unclear why this single mutation has such a profound effect on virulence.

In addition to its natural evolution, the spike protein has been engineered extensively to create more effective vaccine constructs and soluble versions of the trimer that can be used for *in vitro* experiments. In spike-2P (S-2P), one of the first engineered constructs^15,16^, the transmembrane helices are replaced with a synthetic trimerization domain (T4 fibritin or foldon) and the multibasic furin cleavage site (R-R-A-R) is ablated with various substitutions^15–17^. Additionally, two proline mutations, K986P and V987P (designated as 2P) were introduced at the S1 proximal end of the S2 domain. These proline substitutions were designed to prevent the reorganization of the S2 domain necessary for fusion, stabilizing protein’s prefusion conformation^18^. Similar proline variants were engineered previously to stabilize other class 1 viral fusion proteins, including respiratory syncytial virus^19^ and MERS^18,20^. The engineered S-2P variant has led to crucial structural insights and facilitated the rapid development and deployment of vaccine constructs worldwide. However, a detailed comparison of S-2P with membrane-bound spike variants or spike on infectious virus is lacking.

While studies to date indicate that these changes to spike (natural and engineered) do not alter the overall fold of the spike protein^10,13,14,21^, less is known about how these changes alter the dynamics and energetics of the protein. The conformational dynamics of the spike protein have been interrogated by several techniques such as hydrogen-deuterium exchange monitored by mass spectrometry (HDX-MS)^22–26^, molecular dynamics simulations^27,28^ and single-molecule Förster resonance energy transfer^29^. Previous work has demonstrated that HDX-MS is a particularly powerful tool for mapping the conformational heterogeneity of spike^30^. HDX-MS studies on S-2P constructs as well as the isolated receptor binding domain (RBD), have also determined antibody epitopes and evaluated the effect of receptor binding on the rest of the spike protein^22–26^.

In our previous HDX-MS studies on the spike protein, we discovered that S-2P D614 undergoes a very slow (t_1/2_ of hours) conformational change, wherein most of the trimer interface is lost while each protomer remains in its native fold^30^. This conformational change is reversible and temperature-dependent, such that incubation at 4 °C promotes the formation of the ‘open-interface trimer’, while incubation at 37 °C returns the population to the canonical ‘closed-interface trimer’ prefusion conformation. It is important to note that this is different from the well-known conformational change where the RBDs can exist in an ‘up’ or ‘down’ state. To our knowledge, all previous HDX-MS studies on full-length spike, including our prior experiments, have been limited to S-2P and other engineered soluble constructs.

Here we interrogate the conformational dynamics of spike protein in its native membrane environment by carrying out HDX on engineered virus-like particles displaying the spike protein. This native-like environment allows us to evaluate the effects of the multibasic furin site, the engineered stabilizing prolines, the membrane and other changes. We find the open-interface trimer is not unique to soluble S-2P and is also sampled on virus-like particles. Furthermore, the presence and resulting cleavage of the multibasic furin site leads to an allosteric destabilization of the S2’ protease cleavage site as well as increases the proportion of open-interface trimer, which is compensated for by the D614G mutation. Importantly, we also find that the trimerization domain and stabilizing prolines have minimal effect on the fold and conformational landscape of the SARS-CoV-2 spike protein making them a faithful proxy for the membrane-bound, unstabilized spike protein.

## Results

### Hydrogen deuterium exchange of spike protein on enveloped virus-like particles

To study the spike protein in a more native-like environment, enveloped virus-like particles (eVLPs) were made as previously described by genetically appending an endocytosis prevention motif (EPM) and an ESCRT and ALIX-binding region (EABR) to the intracellular, C-terminal end of the full-length spike protein including the transmembrane helices (residues 1-2052)^31^. This leads to the formation of extracellular vesicles that are densely coated with spike protein and are non-infectious and non-replicative. Thus these spike-eVLPs are compatible with HDX-MS methods without the safety concerns of using pseudovirus or live SARS-CoV-2. Modifications were made to our previous spike protein HDX-MS protocol in order to remove the phospholipids in the eVLPs to prevent clogging the columns in the liquid chromatography system (See Methods). Despite the challenges of doing HDX-MS on a heavily glycosylated membrane protein in a complex mixture of other proteins in the eVLP, we identified and processed 200-250 unique spike peptides observed in every experiment, which results in an overlapping 50-55% sequence coverage of ectodomain between S-2P and the spike-eVLP constructs at an average redundancy of 2.7-2.8 (Supplemental Table 1). All spike constructs evaluated in this study have HDX-MS uptake profiles consistent with soluble S-2P and the expected overall structure of prefusion spike (Supplemental Figure 1). Furthermore, we do not observe any bimodal exchange that would suggest the presence of a significant population of post-fusion trimer.

### Unstabilized (K986, V987) D614G SARS-CoV-2 spike with a multibasic furin site samples the open-interface trimer on eVLPs

Our previous studies using solubilized spike (S-2P and its derivatives) revealed that, in addition to the canonical prefusion conformation, the spike protein samples an open-interface trimer conformation. This alternative conformation was detected by bimodal HDX spectra, where the less exchanged distribution corresponds to the prefusion conformation, and the more exchanged distribution corresponds to the open-interface trimer (Figure 1A). The relative populations of each conformer can be determined by fitting the bimodal spectra to two distributions.

**Figure 1.**
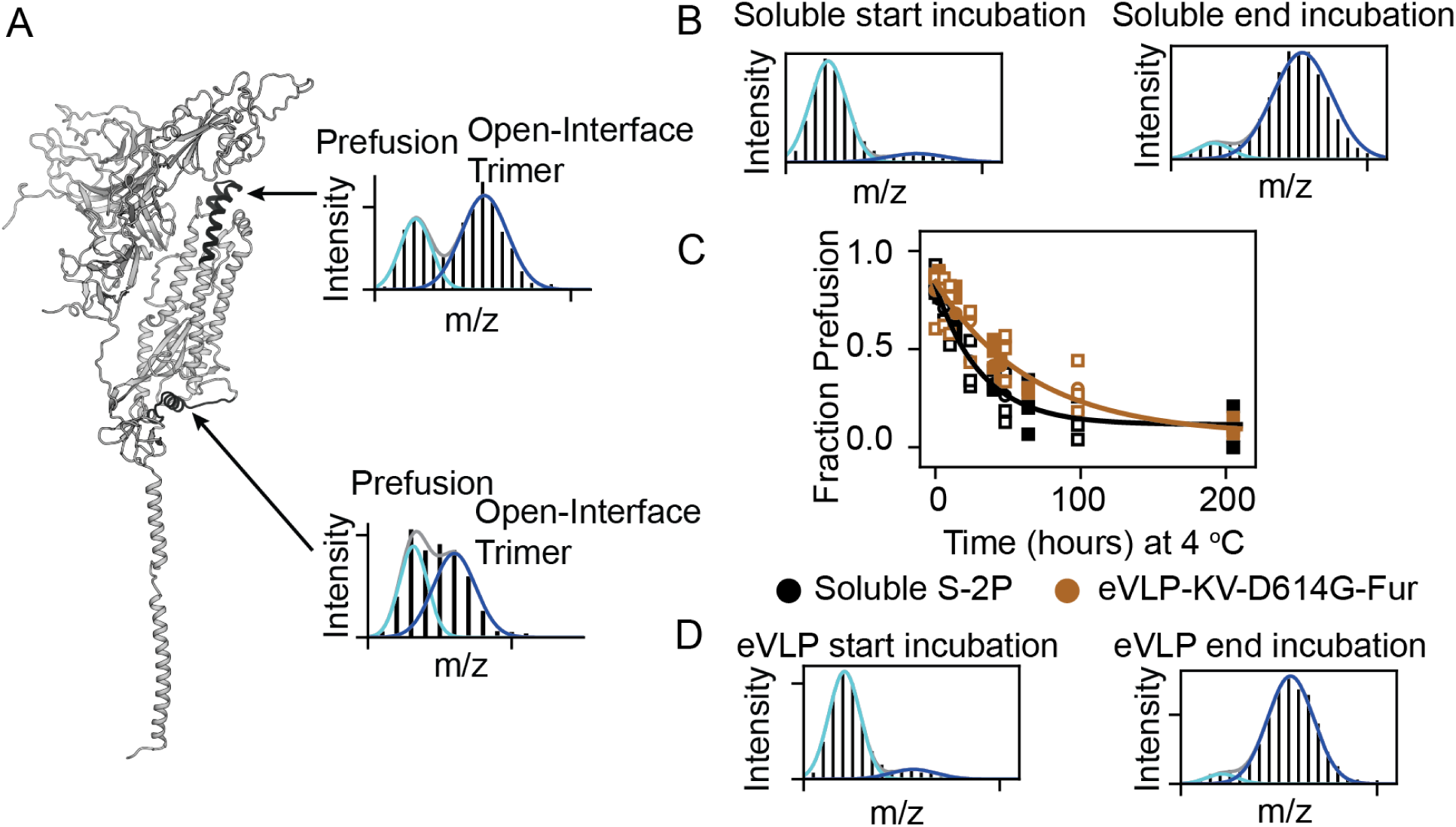
Monitoring the prefusion vs open-interface trimer conformational change. (A) Spike monomer model (full length spike model^28^, black regions are reporter regions used to determine conformation as they exchange differently in the two conformations and show bimodal behavior. Example spectra from the eVLP-S-KV-D614G-Fur from the top and bottom of S2 after 45 hours of incubation at 4 °C. The proportion of prefusion (cyan curve) and open-interface trimer (blue curve) is about the same for both regions. (B) Example spectra from the beginning (0h) and end (205h) of the soluble S-2P 4 °C incubation. (C) Time course of interconversion to open-interface trimer at 4 °C for soluble S-2P and eVLP-S-2P-D614-ΔFur (D) Example spectra from the beginning (0h) and end (205h) of the eVLP-S-KV-D614G-Fur 4 °C incubation.

To address whether the wild-type SARS-CoV-2 spike sequence samples the open-interface trimer conformation on a membrane, eVLPs displaying spike without stabilizing prolines (K986 and V987), with D614G and a cleavable multibasic furin site (R-R-A-R) (eVLP-S-KV-D614G-Fur) were expressed and purified for HDX-MS studies (See Methods). Spike displayed on eVLPs with a multibasic furin site are nearly 100% cleavage at the S1/S2 furin site^31^. We find that the open-interface trimer conformation is indeed sampled by this wild-type SARS-CoV-2 spike sequence on a membrane (Figure 1). We see bimodal spectra at the regions previously shown to be reporters of the two conformations in S-2P (Figure 1B-D), which we can use to quantify the percent of each conformation (See Methods). After incubating the purified spike-eVLPs for 20 hours at 37 °C the eVLP-S-KV-D614G-Fur spike protein is 83 ± 9% closed trimer conformation (Table 1) while for S-2P the 37 °C steady-state closed trimer population is 84 ± 4%. As seen with soluble S-2P, incubation at 4 °C shifts the equilibrium to the open-interface trimer. The rates of interconversion are of the same order of magnitude (Figure 1C), with a halftime of interconversion for eVLP-S-KV-D614G-Fur spike protein of 44 ± 6.8 hours, while under the same conditions the halftime of interconversion for S-2P is 23.7 ± 2.7 hours (Table 1). Therefore, the native spike sequence and environment slows the conversion to the open-interface trimer compared to S-2P while the equilibrium populations are relatively similar.

**Table 1.**

| Construct | % Prefusion after 20 hours of incubation at 37 °C | Rate of opening at 4 °C (h<sup>-1</sup>) | Halftime of opening at 4 °C (h) |
| --- | --- | --- | --- |
| Soluble S-2P | 84 ± 4 | 0.03 ± 0.003 | 23.7 ± 2.7 |
| eVLP-KV-D614G-Fur | 83 ± 9 | 0.02 ± 0.002 | 44.1 ± 6.8 |
| eVLP-KV-D614G-ΔFur | 89 ± 6 | 0.006 ± 0.003 | 117.5 ± 51.2 |
| eVLP-KV-D614-Fur | 50 ± 5 | 0.04 ± 0.009 | 17.3 ± 4.0 |
| eVLP-KV-D614-ΔFur | 89 ± 8 | 0.01 ± 0.002 | 58.0 ± 13.7 |
| eVLP-2P-D614-ΔFur | 95 ± 4 | 0.04 ± 0.004 | 16.7 ± 1.5 |
| eVLP-2P-D614-Fur | 28 ± 7 | 0.06 ± 0.02 | 12.2 ± 3.9 |

### Cleavage at the furin site leads to increased flexibility at the S2’ site

To determine the effects of furin cleavage, we carried out hydrogen exchange on eVLPs displaying spike with K986/V987 and D614G with the multibasic furin site as well as the matched construct where the furin site is replaced by a single alanine (eVLP-S-KV-D614G-ΔFur). With the notable exception of residues 783-822, the majority of the protein shows almost no difference in deuterium uptake between eVLP-S-KV-D614G-Fur and eVLP-S-KV-D614G-ΔFur. For peptides spanning residues 783-822, exchange is significantly greater in the variant with the cleaved multibasic furin site (see Methods, Figure 2A-C). This region contains the fusion peptide as well as the S2’ site, which is cleaved by TMPRSS2 to initiate fusion. The notable increase in hydrogen exchange indicates that the region containing the S2’ cleavage site becomes more flexible when the S1/S2 junction is cleaved. Because the serine protease TMPRSS2 requires an unstructured region for efficient cleavage^32^, processing at the multibasic furin site should increase TMPRSS2 activity at the S2’ site, a potentially important allosteric effect.

**Figure 2.**
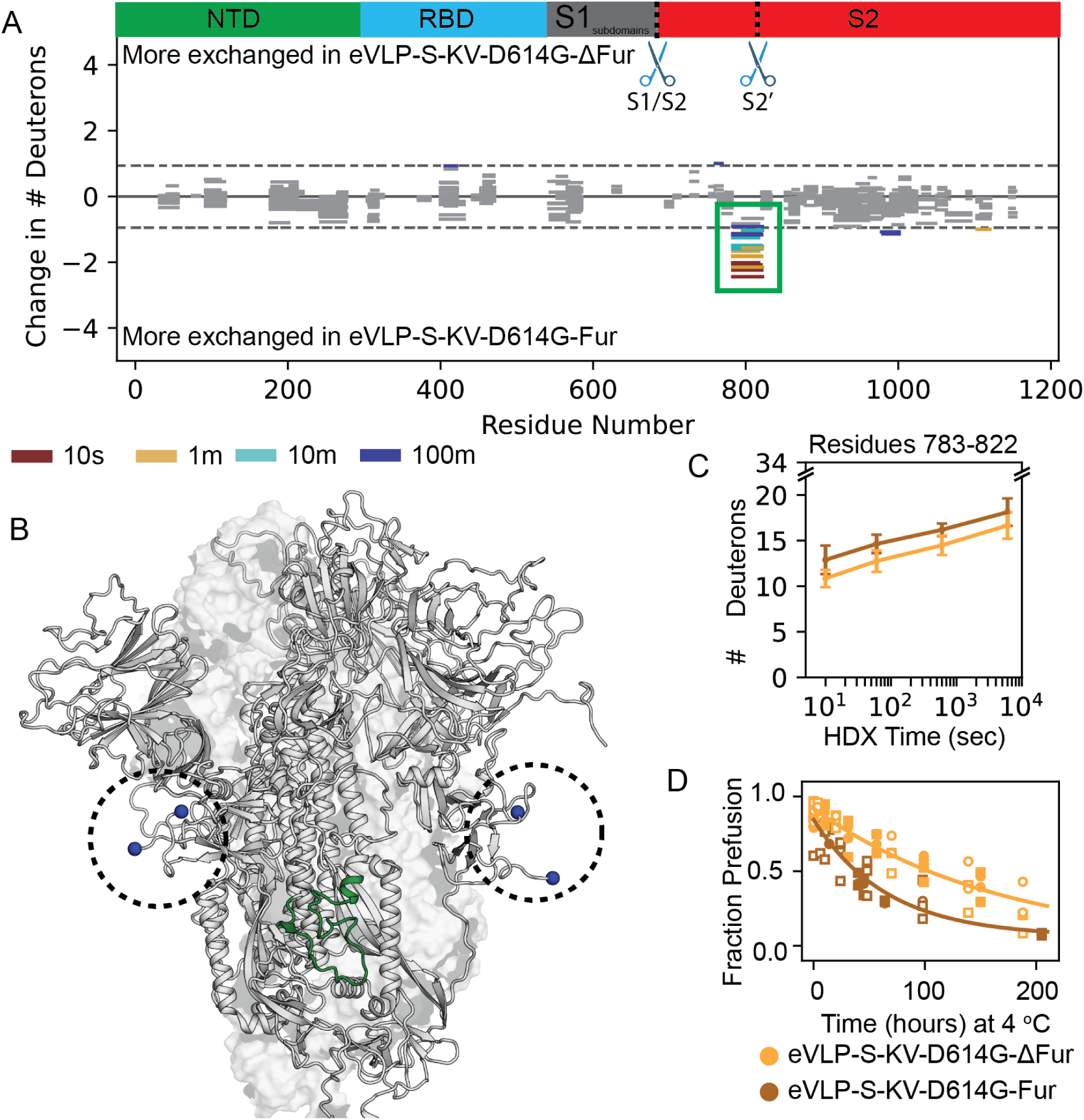
Comparison of eVLP-S-KV-D614G-Fur and eVLP-S-KV-D614G-ΔFur. (A) Woods plot of hydrogen exchange comparing eVLP-S-KV-D614G-Fur and eVLP-S-KV-D614G-ΔFur. Each line is a peptide, the x-axis position represents the sequence it covers and the y-axis position is the difference in exchange. The color of the line encodes the time of exchange. The green rectangle highlights the region with a significant difference in exchange at all timepoints. The exchange significance threshold was determined by the average variability of exchange within replicates of the same system (Supplemental Table 2) and is represented by the grayed out peptides, which are below the threshold. (B) Structural model of spike^28^ with two chains rendered as cartoons and one chain rendered as a white surface. The peptide showing a significant difference in exchange from part (A) is colored green. The location of the S1/S2 furin cleavage site is circled and colored in blue spheres. (C) Uptake plot for the peptide designated in A. Error bars are the standard deviation of 3 replicates. (D) Kinetics of conformational interconversion at 4 °C for eVLP-S-KV-D614G-Fur and eVLP-S-KV-D614G-ΔFur.

The presence or absence of the furin cleavage site does not appear to have a notable effect on the open-closed trimer interface conformational change in the KV D614G background. After 20 hours of incubation at 37 °C the steady-state populations of closed-interface, or prefusion trimer is 89 ± 6% for eVLP-S-KV-D614G-ΔFur and 83 ± 9% for eVLP-S-KV-D614G-Fur. The presence of the furin site does increase the rate of opening to the open-interface trimer. The t_1/2_ of opening at 4 °C is 117.5 ± 51.2 hours for eVLP-S-KV-D614G-ΔFur and only 44.1 ± 6.8 hours for eVLP-S-KV-D614G-Fur (Figure 2D, Table 1).

### G614 stabilizes the closed-interface trimer conformation compared to D614 in the presence of the cleavage at the S1/S2 furin site

We next investigated the role of the D614G mutation by comparing spike-eVLP constructs with D614 to spike-eVLPs with G614. Position 614 is in the S1 domain facing the S2 domain (Figure 3A). When the multibasic furin site is cleaved, a glycine at position 614 dramatically stabilizes the prefusion conformation compared to the same construct with aspartic acid at position 614 (Figure 3B). After 20 hours of incubation at 37 °C, eVLP-S-KV-D614G-Fur is 83 ± 9% in the prefusion conformation (closed-interface trimer) compared to eVLP-S-KV-D614-Fur, which is only 50 ± 5% prefusion (Table 1). This effect, however, is highly dependent on the presence of the furin cleavage; in the absence of the furin site, D614G does not have an effect on the equilibrium percentage of prefusion conformation (89 ± 8% for eVLP-S-KV-D614-ΔFur compared to 90 ± 6% in eVLP-S-KV-D614G-ΔFur). Thus G614 stabilizes the closed interface trimer in comparison to constructs with D614, but only in the presence of the cleaved S1/S2 furin site. D614G also slows the trimer interface opening approximately two-fold in the presence and absence of the multibasic furin site (Figure 3B, Table 1).

**Figure 3.**
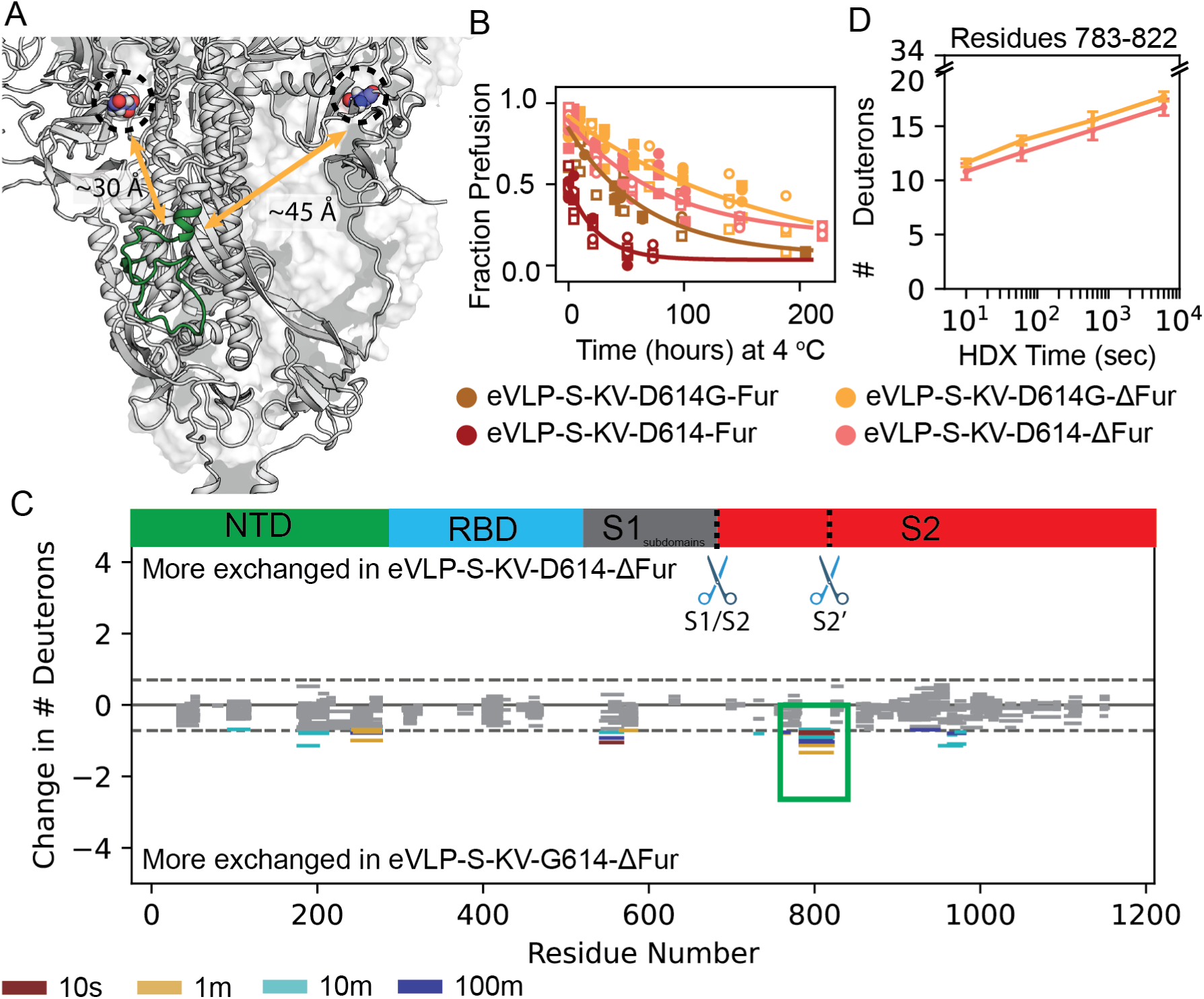
Comparison of eVLP-S-KV-D614-ΔFur and eVLP-S-KV-D614G-ΔFur. (A) Model of spike^28^ with two chains rendered as cartoons and one rendered as a white surface. The peptide showing a significant difference in exchange from part (C) is colored in green. The location of D614 is shown on the same monomer and the adjacent monomer in blue spheres and circled. The distance between the peptide containing the S2’ cut site and the D614G mutation on the same protomer and the adjacent protomer is indicated with the orange arrows. (B) Kinetics of conformational interconversion at 4 °C for eVLP-S-KV-D614-ΔFur, eVLP-S-KV-D614G-ΔFur, eVLP-S-KV-D614-Fur and eVLP-S-KV-D614G-Fur. (C) Woods plot of hydrogen exchange comparing eVLP-S-KV-D614-ΔFur and eVLP-S-KV-D614G-ΔFur. Each line is a peptide, the x-axis position represents the sequence it covers and the y-axis position is the difference in exchange. The color of the line encodes the time of exchange. The green rectangle highlights the region showing significant differences in exchange. The exchange significance threshold was determined by the average variability of exchange within replicates of the same system (Supplemental Table 2) and is represented by the grayed out peptides, which are below the threshold. (D) Uptake plot for the peptide designated in C. Error bars are the standard deviation of 3 replicates.

When using hydrogen exchange rates to compare changes in local flexibility, it is important to compare samples populating the same conformational distribution, preferably predominantly in one or the other conformation. Otherwise the differences may be attributable to the different conformations rather than changes in flexibility. In the presence of furin cleavage, the change from D614 and D614G has a large effect on the conformational equilibrium (After 20 hours of incubation at 37 °C eVLP-S-KV-D614-Fur is only 50% prefusion, whereas eVLP-S-KV-D614G-Fur is 83% prefusion. We can, however, directly compare D614 and D614G in the background of the ablated furin site, (comparing eVLP-S-KV-D614-ΔFur to eVLP-S-KV-D614G-ΔFur) as both are approximately 89% prefusion (Table 1). In continuous hydrogen exchange, only one region shows a significant difference - the region of the S2’ cleavage site and fusion peptide (Figure 3C-D) - with more exchange observed in G614 compared to D614. Thus the D614G mutation results in an increased flexibility of the S2’ protease site in the absence of furin cleavage.

### Spike samples the open-interface trimer in the presence and absence of the stabilizing prolines

The two proline mutations (K986P and V987P) are a feature of the soluble S-2P and all its variants. Residues 986 and 987 reside in a loop on the spike protein between HR1 and the central helix. In addition to increasing the expression of soluble, stable spike variants, they were designed to prevent spontaneous, irreversible adoption of the post-fusion conformation ^16,18^. Licensed mRNA vaccines for COVID-19 include the two proline mutations with the multibasic furin cleavage site ^4^. To evaluate the effect of these prolines on both the formation of the open-interface trimer as well as the overall structure of the protein we compared eVLPs displaying spike with D614 and either the native KV residues or the proline substitutions with and without furin cleavage (eVLP-S-KV-D614-Fur vs eVLP-S-2P-D614-Fur and eVLP-S-KV-D614-ΔFur vs eVLP-S-2P-D614-ΔFur). In the absence of furin cleavage, the two proline substitutions have small effects on the steady-state populations at 37 °C and slightly increase the trimer interface opening rate (Figure 4A, Table 1). In the presence of furin cleavage, the proline substitutions lower the steady-state prefusion population at 37 °C even more dramatically to 28% compared to 50% for a construct with native residues at these sites (Table 1). Thus, this conformational change is not a unique feature of and is not dramatically affected by the engineered proline substitutions.

**Figure 4.**
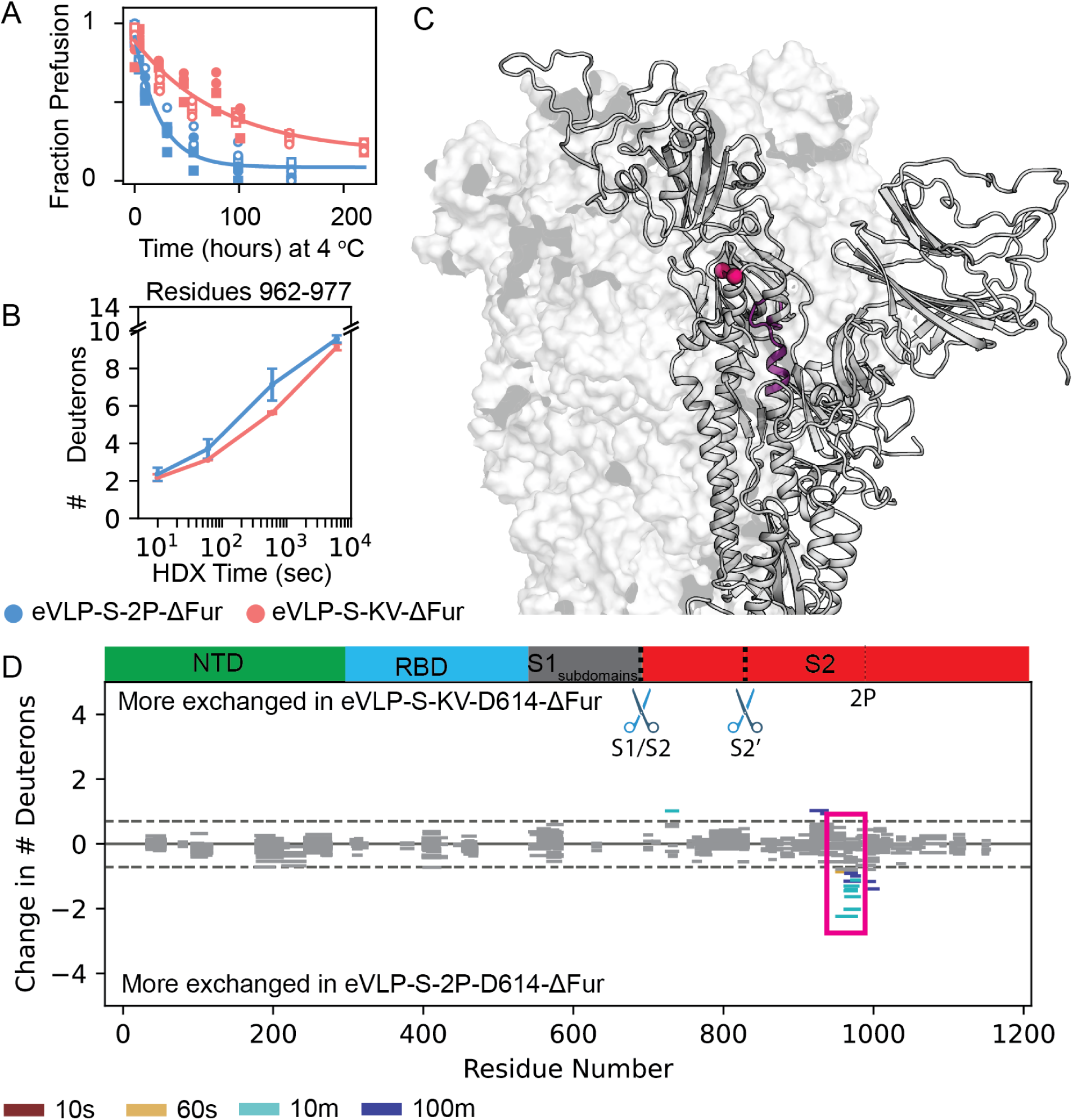
Comparing stabilizing prolines to native K986 and V987. (A) Time course of conformation interconversion at 4 °C for eVLP-S-2P-D614-ΔFur and eVLP-S-KV-D614-ΔFur. (B) Uptake plot of a peptide covering residues 961-977. Error bars are the standard deviation of 3 replicates. (C) The location of the peptide (purple) from (B) on a model of spike^28^. Location of K986/V987 is shown in red. One chain rendered as a cartoon, two are rendered as a surface. (D) Woods plot of hydrogen exchange comparing eVLP-S-2P-D614-ΔFur and eVLP-S-KV-D614-ΔFur. Each line is a peptide, the x-axis position represents the sequence it covers and the y-axis position is the difference in exchange. The color of the line encodes the time of exchange. Region of difference from part (B) in magenta rectangle. The exchange significance threshold was determined by the average variability of exchange within replicates of the comparison (Supplemental Table 2) and is represented by the grayed out peptides, which are below the threshold.

Most structural studies of SARS-CoV-2 spike use soluble ectodomains containing two proline substitutions and thus there is relatively little information about the structural consequences of these proline substitutions. A cryo-EM study on detergent solubilized full-length spike with the native K986/V987 residues showed the S1 subunit interfaces are more tightly packed than in structures of soluble S-2P^21^. They suggest this is due to the absence of a salt bridge between K986 and D427 or D428 after proline mutation of K986, resulting in unsatisfied charges leading to a looser structure. This is consistent with continuous labeling experiments on eVLP-S-KV-D614-ΔFur and eVLP-S-2P-D614-ΔFur, where we see no large-scale changes to the overall structure and flexibility of the protein (Figure 4B-D). We do, however, see a small yet significant difference in exchange behavior in regions covering the C-terminal end of HR1 and the loop before the central helix (Figure 4B-D), with more protection in the KV variant. Since this region is adjacent to the site of the proline substitutions (Figure 4C), this is consistent with more flexibility in the S1 domain as seen by cryo-EM^21^.

### The membrane environment slightly stabilizes the spike protein and does not change the fold

Soluble variants of spike, including S-2P, have the transmembrane helices replaced by a T4 fibritin trimerization domain. This trimerization domain preserves the trimeric nature of the protein; however, it is possible that strain induced by this engineered domain biases the formation of the open-interface trimer. To address this question, we compared soluble S-2P to a sequence-matched eVLP construct (eVLP-S-2P-D614-ΔFur). After incubating at 37 °C for 20 hours the eVLP-S-2P-D614-ΔFur is approximately 95 ± 4% prefusion conformation compared to 84 ± 4% for the soluble S-2P construct (Figure 5A, Table 1). Thus, the membrane may slightly stabilize the closed-interface trimer conformation relative to the soluble matched construct. Under these conditions, the two proteins have similar opening rates at 4 °C (Table 1, Figure 5A).

**Figure 5.**
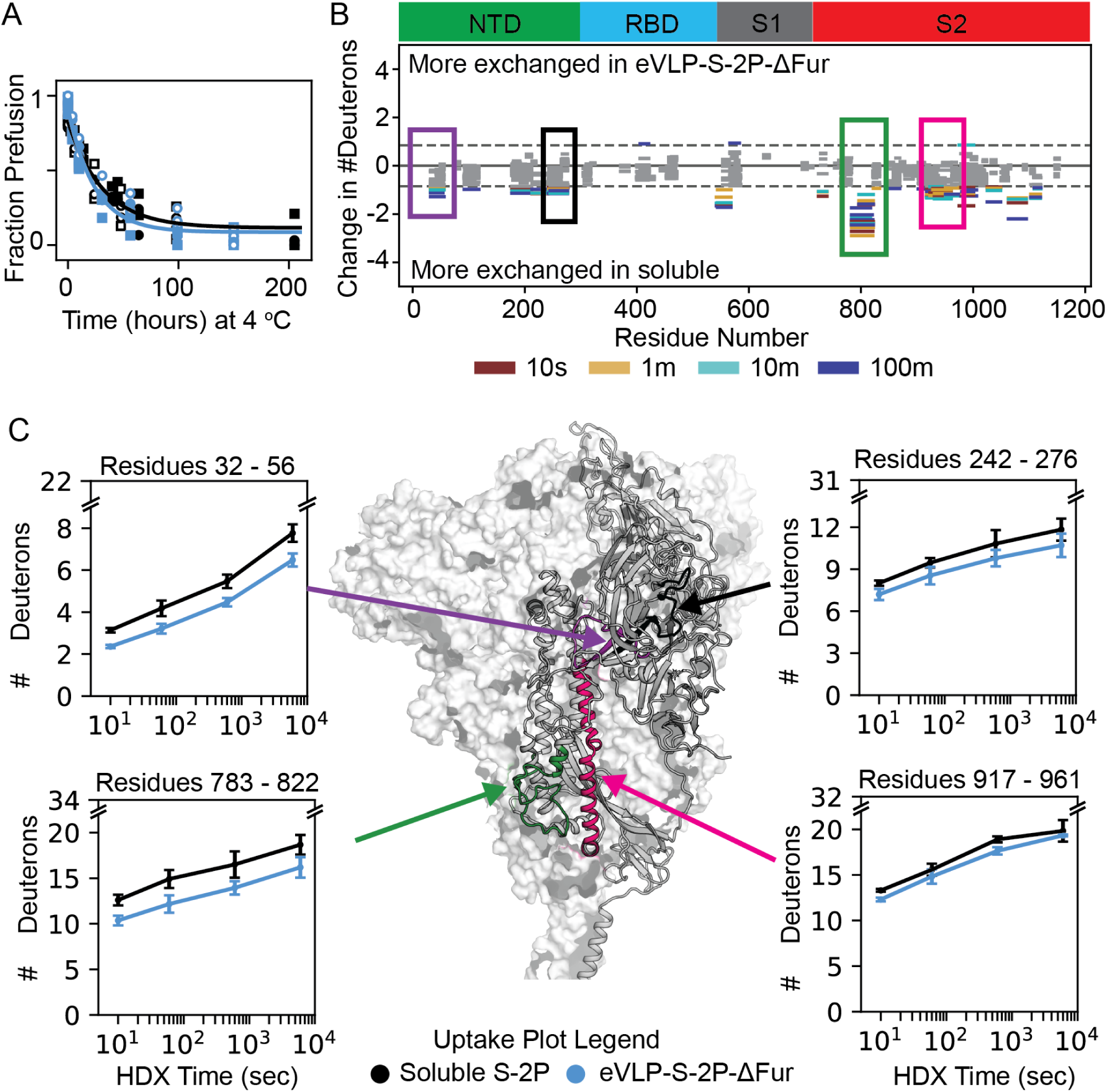
Comparison of soluble S-2P and eVLP-S-2P-D614-ΔFur. (A) Kinetics of conversion to the open-interface trimer at 4 °C of soluble S-2P and eVLP-S-2P-D614-ΔFur spike. (B) Woods plot comparing deuterium uptake between soluble S-2P and eVLP-S-2P-D614-ΔFur. Each line is a peptide whose x-axis value and length represents its sequence position and the y-axis value is the difference in the exchange. The color of the line represents the deuteration time. The exchange significance threshold was determined by the average variability of exchange within replicates of the same comparison (Supplemental Table 2) and is represented by the grayed out peptides, which are below the threshold. The boxed peptides are selected for illustrative purposes and the corresponding uptake plot is shown in parts (C) and the location of these peptides is shown on a model of the spike structure^28^ with one chain rendered as a cartoon and two chains rendered as a white surface. Error bars on the uptake plots are the standard deviation of 3 replicates.

We conducted continuous labeling HDX on S-2P and the sequence matched spike displayed on an eVLP to determine if the T4 fibritin trimerization domain and resulting solubilization alters the hydrogen exchange pattern of the spike protein compared to the more native membrane environment. While the hydrogen exchange uptake patterns are similar between the two constructs in general, we see more exchange in soluble S-2P throughout most of the protein (Figure 5B). This suggests that eVLP-S-2P-D614-ΔFur is globally more stable than soluble S-2P, which is supported by the increased exchange along the central helix. Overall, the uptake patterns between soluble S-2P and eVLP-S-2P-D614-ΔFur suggests that the two constructs have similar folds with small differences in local stability throughout the protein (Figure 5C).

## Discussion

By combining HDX-MS with the ability of eVLPs to display full-length native-like SARS-CoV-2 spike protein, we have determined the energetic and conformational effects of both the membrane environment and unique sequence features that are considered incompatible with soluble protein constructs. We demonstrate that the SARS-CoV-2 spike protein ubiquitously samples both the closed-interface canonical prefusion conformation and open-interface trimer conformation in the native membrane across all constructs examined. The relative proportions and the energy barrier to opening vary based on sequence modifications. The well-studied soluble constructs also contain two proline substitutions that do not appear to greatly alter the protein’s energy landscape. Remarkably, we find that prohibiting furin cleavage and the early emergence of D614G, induce global, local, and allosteric changes in protein stability, offering valuable insights into protein design and viral evolution.

### Implications for viral evolution

We find that cleavage at the furin site results in a significant increase in local flexibility around the S2’ protease site and the fusion peptide region. This change in dynamics is distant (30 Å away) from the furin-cleavage site and therefore is not simply a result of an increase in solvent accessibility arising from the cleavage of the S1/S2 junction. The observed disordering of the S2’ site may explain the noted evolutionary advantage of the multibasic furin cleavage site. A more disordered S2’ site should be more efficiently processed by TMPRSS2, a serine protease, which requires a disordered stretch of amino acids in order to bind and cleave. More efficient cleavage at the S2’ site should lead to more efficient membrane fusion. While the structures of S1/S2 cleaved soluble spike protein have been reported, the only noted changes are modest reorientation of S1 subdomains^13^ as well as differences in packing along the interface between the S1 domain and S2 domain^21^, but no changes at the S2’ site were noted. To our knowledge, our HDX study provides the first molecular explanation for why furin cleavage is advantageous to the virus.

Our results also shed light on the effects of the early-adopted substitution at position 614 (D614G). While structural studies report no large-scale structural changes, some note small changes such as rotations of S1 domains^10,13^, an increase in secondary structure of the 640 loop in S1^14^, as well as changes in the RBD up-down equilibrium^13^. Our HDX-studies also suggest that D614G does not significantly alter the structure of the SARS-CoV-2 spike protein. We do, however, see some small, yet statistically significant, changes in deuterium uptake in S1 and HR1 that could reflect the local changes noted in the structural studies^10,14^. In combination, the cryo-EM studies, which would note any large changes in structure, and our HDX results, which report on small structural changes, indicate that the advantage conferred by D614G is not due to a dramatic structural change in the spike protein.

Our data do suggest that the D614G mutation stabilizes the canonical prefusion trimer conformation relative to the open-interface trimer particularly in furin-cleaved constructs. Although the opening rates we measure are very slow and unlikely to directly reflect conformational changes during the infection process, they do suggest that D614G stabilizes the trimer interface. Less interface contacts in D614 (compared to D614G) may increase the rate of S1 shedding, since in order for the S1 domain to dissociate from the S2 domain, both intra- and inter-protomer contacts must be lost. While S1 shedding is required for infection to proceed after receptor binding, premature S1 shedding could render the virus non-infectious. Therefore, reducing premature shedding of S1 may be the mechanistic basis for the competitive advantage of the D614G mutation.

In combination, the effects we observe for D614G and the furin cleavage site suggest an interesting evolutionary path. Evolution of the multibasic furin cleavage site at S1/S2, facilitated easier processing of the S2’ site by TMPRSS2; however, this also reduces trimer stability, increasing premature S1 shedding. In response to this premature S1 shedding, the D614G mutation stabilizes the trimer, resulting in a more infectious virus (Figure 6).

**Figure 6.**
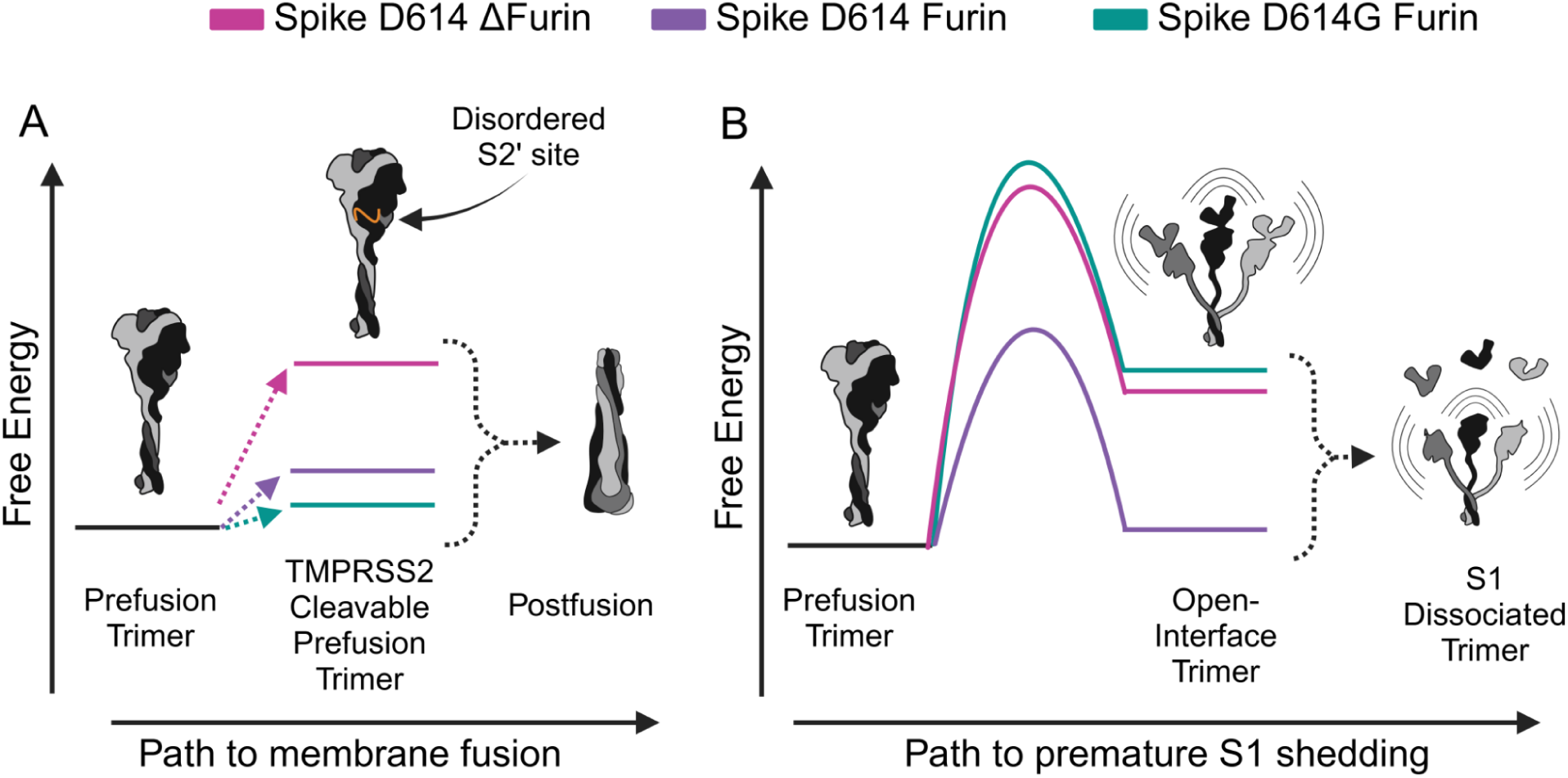
The energetic landscape of the SARS-CoV-2 spike protein and its evolutionary adaptations. In our paper we find that mutations in the SARS-CoV-2 spike protein affect two competing pathways: (A) accessing the TMPRSS2 cleavable state (orange loop) which leads to membrane fusion and (B) formation of the open trimer which may lead to premature S1 shedding. SARS-CoV-2 spike protein with D614 and no furin cleavage site (magenta) minimally populates the open-interface trimer as it is high energy. However, the TMPRSS2-cleavable state is also high-energy as the S2’ site is more ordered. This combination prevents premature S1 shedding but also has low fusion efficiency. When SARS-CoV-2 D614 spike acquired the furin cleavage site (purple), this lowered the energy of the TMPRSS2-cleavable state, enhancing fusion efficiency. However, this adaptation also destabilized the prefusion trimer by lowering the energy of the open-interface trimer, increasing premature S1 shedding and the proportion of non-infectious virus. The subsequent D614G mutation (teal) restores trimer stability by increasing the free energy of the open-interface trimer, leading to reduced S1 shedding while maintaining the low-energy TMPRSS2-cleavable state, thus optimizing viral infectivity. Barrier and free energy heights are to summarize findings from HDX about relative trends in populations and interconversion rates and are for illustrative purposes only.

### Implications for engineering and therapeutics

Our spike-eVLP studies allow us to evaluate if the engineered changes in S-2P affect the flexibility and stability of the native spike. Reassuringly, we find that the overall fold and structure of the spike protein is not altered in going from the ‘native’ membrane bound construct to the soluble S-2P: the deuterium uptake patterns between these two constructs are similar throughout the entire protein. Instead, our data suggest that some of the engineered mutations show more specific, and often local, effects. Replacing the transmembrane helices with a trimerization domain appears to mildly destabilize the entire protein suggesting strain on the geometry of the trimer. The 2P substitution (K986P/V987P) has a more local effect: it destabilizes the C-terminus of the HR1 helix, which is proximal to the mutation, but does not seem to affect the overall fold or global stability. As discussed above, the changes at position 614 (D614/D614G) and the presence or absence of a furin cleavage show interesting allosteric effects. Comparing constructs with and without furin cleavage, we find local changes in uptake in the S2’ site/fusion-peptide region, suggesting these mutations allosterically alter the flexibility of that specific area.

Importantly, our hydrogen exchange studies provide important insights into the impact of these engineered changes on the propensity for the spike protein to adopt the open-interface trimer. The propensity to open and expose the trimeric interface to solvent is inherent to the protein and is not a consequence of the solubilization or proline mutations. In fact, the only engineered change that seems to have a dramatic effect is the removal of the native multibasic furin cleavage site. Thus, we conclude that the open-trimer conformation is on the energy landscape and therefore sampled on a bonafide SARS-CoV-2 virus.

While our studies identify the structural and dynamic effects of these engineered changes, it is difficult to deduce how these changes affect the antigenicity of vaccine constructs or the affinity of therapeutic antibodies. Small changes in flexibility may alter the affinity of an antibody raised against constructs biased towards one conformation or another, but these effects are hard to predict. It is worth noting, however, that most of the differences we see are not in the NTD and RBD, which are considered the most antigenically important regions of the SARS-CoV-2 spike protein. Neutralizing antibodies have been identified that target the fusion peptide of the SARS-CoV-2 spike protein^33^, an area where we do see a difference in exchange between constructs with and without furin cleavage and with D vs G as position 614. Given the increase in flexibility of this region in the background of furin cleavage at the S1/S2 junction, the affinity of antibodies could be different in constructs with an ablated furin site compared to ones with a cleavable furin site. Interestingly, in order to measure the affinity of fusion peptide-targeting antibodies, these authors used an S-2P construct with a cleavable furin site, suggesting that to measure tight binding furin cleavage was necessary. Furthermore, binding was much tighter to an isolated S2 construct, suggesting that an increased dynamics in this region leads to tighter binding. This is one example of how a change in dynamics can affect antibody binding and thus is an important consideration when evaluating affinity.

In sum, our ability to harness recently developed eVLP technology for HDX-MS studies has opened the door to studies of viral fusion proteins in environments that closely mimic their native state. This approach has provided critical structural insights into viral evolution and infectivity while also illuminating how protein engineering efforts result in modifications of the local dynamics of spike proteins. The potential of HDX-MS paired with eVLPs to revolutionize the study of other viral fusion proteins and membrane proteins in general is immense. Membrane protein research has long been hampered by the challenges of expression, purification, and the use of detergents, which can distort their conformational landscapes compared to native lipid environments^34–36^. By embedding proteins within eVLPs, we achieve a setting that closely resembles the natural viral membrane. Moreover, eVLPs offer substantial safety advantages due to their lack of genetic material and enable significantly higher protein concentrations than alternative methods like pseudoviruses. These benefits position eVLPs as an excellent platform for HDX-MS studies. Looking ahead, the integration of eVLPs with HDX-MS holds enormous potential for deepening our understanding of viral fusion proteins and other membrane proteins, shedding light on the structural and energetic impacts of evolutionary and engineered mutations.

## Methods

### Protein expression and purification

SARS-CoV-2 Spike (S-2P) was expressed and purified from stably transformed Expi293 cells, following methods as described^37^. Briefly, stable Expi293 suspension cells were maintained in Expi293 media at 37 °C. Cells were grown for 6 days, then harvested by centrifugation, and filtered. Supernatant was adjusted to pH 7.4 and loaded onto a 5 mL HisTrap Excel column and washed with 60 CV of buffer. Captured proteins were eluted with 10 CV of (20 mM sodium phosphate, 500 mM NaCl, 500 mM imidazole, pH 7.4). Eluted proteins were buffer exchanged into PBS using 100 kDa MWCO Amicon concentrators and filtered prior to storage at −80°C.

### Spike EABR-eVLP Cloning, Expression and Purification

Full length SARS-CoV-2 spike protein constructs were cloned into a p3BNC plasmid containing the ALIX and EABR tags as previously described^31^. Mutations (D614/D614G, ΔFur) were introduced via “around the horn” PCR. The ΔFur mutation was made by replacing the R-R-A-R residues with a single alanine^17^. The resulting plasmid was transformed into MachOne E.coli RbCl_2_ chemically competent cells via heatshock at 42 °C and plated onto ampicillin LB plates. A 100 mL overnight culture was then inoculated with a colony from the selection plate or a stab from a glycerol stock from a previous overnight culture. The culture was then prepped with a Qiagen maxi-prep kit.

Expi 293 Freestyle cells were prepared in various culture sizes (50-150 mL) at 2.5e6 cells/mL in FreeStyle™ 293 Expression Medium (Gibco, ThermoFisher). To transfect cells, 1 ug DNA per milliliter of cell culture was filtered with 300 uL of OptiMEM media (Gibco, ThermoFisher). The filtered DNA was diluted to 25 ug/mL in OptiMEM. In a separate tube 4 ug of PEI Max (Kyfora Bio) was prepared in 0.4 mL of OptiMEM for every 1 mL of cell culture and mixed by inverting the tube 5-10 times. The DNA mixture was then added to the PEI Max mixture and allowed to complex for 15 minutes before adding dropwise to the cell culture while swirling the flask. The cells were allowed to express for 65 - 70 hours at 37 °C and 8% CO2 before harvesting by spinning down the cells at 4000xg for 15 minutes and collecting the supernatant.

To purify eVLPS, the collected supernatant was passed through a 0.2 um filter and then was set up for sucrose cushion centrifugation. Optiseal 29.9 mL ultracentrifuge tubes (Beckman Coulter) were filled with 25 mL of filtered supernatant and then 4.9 mL of 20% sucrose PBS was layered at the bottom of the tube. The tubes were balanced, capped and then spun in an MLA50 rotor in a 10 °C Optima MAX-XP for 2 hours at 48,000 rpm (rcf average = 150,700). The tubes were then emptied and the tubes were cut in half at an angle. 250 uL of 1X PBS was added to the pellet and then the cut tube was covered with parafilm. The pellet was then allowed to incubate overnight at room temperature before pipetting up and down to resuspend. The resuspended pellets of the same construct were combined and incubated at 37 °C for 15 minutes in an 1.5 ml eppendorf tube. The tubes were then spun down at 11,000xg in a tabletop centrifuge for 2 minutes and the supernatant was collected. This was repeated 4 times to remove insoluble aggregates. The final supernatant was then flowed over an S200 sizing column to separate the eVLPS (eluting in the void volume) from soluble protein (eluting later from the column). The void fraction was then pooled and concentrated to 20 uL for every 25 mL of culture. The eVLPs were then either used immediately for experiments or were flash frozen in liquid nitrogen and stored at −80 °C.

### Continuous Labelling HDX-MS

To initiate exchange, concentrated eVLPs or 1.67 uM soluble S-2P spike trimer was diluted 10-fold to 1X PBS deuterated buffer (made by lyophilizing 1X PBS (Sigma-Aldrich P4417) and resuspending the powder with D_2_O (TCI-America Cat No: W0002)) at pH_read_ 7.4. At each timepoint, exchange was quenched by mixing 100 uL of exchange reaction with 100 uL of ice-cold 2X offline digestion quench buffer containing zirconium oxide (3M Urea 20 mM TCEP pH 2.4 with 25 mg/ml Zirconium oxide) and adding 2 uL of a mixture of aspergillopepsin (Sigma-Aldrich P2143) and porcine pepsin (Sigma-Aldrich P6887) (3.5 mg/mL aspergillopepsin, 5 mg/mL porcine pepsin). The mixture was vortexed for 3 seconds before incubating on ice for 57 seconds, vortexing for 3 seconds and then incubating on ice for another minute for a total quench time of 2 minutes. The mixture was put into a Costar SpinX 0.22 um filter (Corning P/N: 8160) and spun at 6000 rcf for 30 seconds. The filtered sample was then moved to a clean tube and flash frozen in liquid nitrogen. Samples were stored at −80 °C until LC-MS.

### Incubation kinetics and pulse labeling

For evaluating the temperature dependent kinetics of interconversion, concentrated eVLPs or 1.67 μm soluble spike trimer were incubated at 37°C for 20 hours. Samples were then moved to a temperature-controlled chamber at 4°C and the population of each state was evaluated at the specified time points. To evaluate the relative population of the A and B conformer at each time point, 10 μL of spike sample was removed from the incubation tube and mixed with 90 μL of room temperature deuterated buffer. After a one-minute labeling pulse, ice cold 100 μL of quench buffer (3M Urea 20 mM TCEP pH 2.4 with 25 mg/ml Zirconium oxide) was mixed with 100 μL of labeled protein/eVLP. To digest the protein, 2 uL of a mixture of aspergillopepsin (Sigma-Aldrich P2143) and porcine pepsin (Sigma-Aldrich P6887) (3.5 mg/mL aspergillopepsin, 5 mg/mL porcine pepsin) was added to the mixture. The mixture was vortexed for 3 seconds before incubating on ice for 57 seconds, vortexing for 3 seconds and then incubating on ice for another minute for a total quench time of 2 minutes. The mixture was put into a Costar SpinX 0.22 um filter (Corning P/N: 8160) and spun at 6000 xg for 30 seconds. The filtered sample was then moved to a clean tube and flash frozen in liquid nitrogen. Samples were stored at −80 °C until LC-MS.

### Liquid chromatography - mass spectrometry

LC-MS was conducted based on previously reported optimized methods^30^ with a different analytical column. To reiterate, all samples were thawed immediately before injection into a cooled valve system (Trajan LEAP) coupled to a LC (Thermo UltiMate 3000). Sample time points were injected in random order. The temperature of the valve chamber, trap column, and analytical column were maintained at 2 °C. Peptides were desalted for 4 minutes on a hand-packed trap column (Thermo Scientific POROS R2 reversed-phase resin 1112906, 1 mm ID x 2 cm, IDEX C-128) at 200 μL/min. Peptides were then separated with a C18 analytical column (ACQUITY UPLC BEH C18 Column, 130Å, 1.7 μm, 1 mm X 50 mm) and a gradient of 5-40% buffer B (100% Acetonitrile, 0.1% Formic Acid) at a flow rate of 40 μL/min over 14 minutes, and then of 40-90% buffer B over 30 seconds. The analytical and trap columns were then sawtooth washed and equilibrated at 5% buffer B prior to the next injection. Protease columns were washed with two injections of 100 μL 1.6 M GdmCl and 0.1% formic acid prior to the next injection. Peptides were eluted directly into a Q Exactive Orbitrap Mass Spectrometer operating in positive mode (resolution 70000, AGC target 3e6, maximum IT 50 ms, scan range 300-1500 m/z). For each spike construct, a tandem mass spectrometry experiment was performed (Full MS settings the same as above, dd-MS2 settings as follows: resolution 17500, AGC target 2e5, maximum IT 100 ms, loop count 10, isolation window 2.0 m/z, NCE 28, charge state 1 and ≥7 excluded, dynamic exclusion of 15 seconds) on undeuterated samples.

### Peptide identification

Byonic (Protein Metrics) was used to identify unmodified and glycosylated peptides in the tandem mass spectrometry data. The sequence of the expressed construct, including signal sequence and trimerization domain, was used as the search library. Sample digestion parameters were set to non-specific. Precursor mass tolerance and fragment mass tolerance was set to 6 and 10 ppm respectively. Variable N-linked glycosylation was allowed, with a library of human N-glycans. Peptide lists (sequence, charge state, and retention time) were exported from Byonic and imported into HDExaminer 3 (Sierra Analytics). When multiple peptide lists were obtained, all were imported and combined in HDExaminer 3.

### HDExaminer3 analysis

Peptide isotope distributions at each exchange time point were fit in HDExaminer 3.2. For glycosylated peptides, only the highest confidence modification was included in the mass spectra search and analysis. Deuteration levels were determined by subtracting mass centroids of non-deuterated peptides from deuterated peptides. Care was taken to adjust the retention time window for each peptide, so that a consistent slice of the elution peak was taken and was the same width in all conditions to avoid deuteration elution bias. Woods plot significance was calculated in HDExaminer by determining the average variation of deuteration amongst three replicates. Peptides discussed in the manuscript all had timepoints that were greater than the calculated significance threshold and were determined to have a p-value < 0.05, as visualized on a volcano plot.

### Bimodal fitting and conformation quantification

The quantification of each conformation was conducted as previously described^30^. To reiterate, peptide mass spectra for bimodal peptides were exported from HDExaminer 3.0. All quantitative analysis of the exported peptide mass spectra was performed using python scripts in Jupyter notebooks. After importing a peptide mass spectra, the m/z range containing all possible deuteration states of the selected peptide was isolated and the find_peaks method from the scipy.signal package was used to identify each isotope in the mass envelope and the height of each peak was used as its intensity. The area of the total mass envelope was normalized to account for run-to-run differences in intensity. The bimodal mass envelopes for all timepoints under the same condition were globally fit to a sum of two gaussians, keeping the center and width of each gaussian constant across all incubation time points. Fitting was done using the curve_fit function from the scipy.optimize package. After fitting, the area under each individual gaussian was determined to approximate the relative population of each state. Two peptides from the S1 proximal side of S2 (residues 878-1010) and two peptides from the the S1 distal side of S2 (residues 878-925) were processed for each timepoint and then all points were plotted against time and fit to a single exponential. The error of the fitted rate was determined from the covariance of the fitted parameters and then was propagated to determine the error on the halftime. To determine the population of prefusion after 20 hours of incubation at 37 °C the prefusion conformation from all four peptides (two from top of S2 and two from bottom of S2) at the zero-hour time point of the 4 °C incubation were averaged together

## Acknowledgments

We would like to thank the entire Marqusee Lab and Dr. Shawn M. Costello for advice and critical reading of the manuscript. SM is a Chan Zuckerberg Biohub Investigator. Figure 6 created with BioRender.com. This work was funded by the National Science Foundation (2322801), the National Institutes of Health (R35GM149319) and an NIH Director’s Early Independence Award (FAIN# DP5OD033362).

---

**Supplemental Table 1.**

| Data Set | eVLP-S-KV-D614G-Fur | eVLP-S-KV-D614G-ΔFur | eVLP-S-KV-D614-ΔFur | eVLP-S-2P-D614-ΔFur | Soluble S-2P |
| --- | --- | --- | --- | --- | --- |
| HDX reaction details | **Deuteration buffer:** 1X PBS pH(read) = 7.0.<br>**Quench Buffer:** 3M Urea, 20 mM TCEP, 25 mg/mL Zirconium Oxide pH 2.4<br>**Digestion Condition:** 2 uL of 5 mg/mL porcine pepsin + 3.5 mg/mL aspergillopepsin. Digest on ice for 2 minutes |  |  |  |  |
| HDX time course (sec) | 10, 60, 600, 6000 |  |  |  |  |
| HDX control samples | None. To control for exchange, compared replicates were exchanged on the same day with the same buffers. Uptake values were used for comparison only |  |  |  |  |
| Back-exchange | Not backexchange corrected. To control for backexchange, compared replicates were exchanged and injected on the same day. |  |  |  |  |
| # of peptides | 222 | 220 | 223 | 234 | 232 |
| Sequence coverage | 58.20% | 57.71% | 58.20% | 58.20% | 58.20% |
| Average peptide length / Redundancy | 15.71 / 2.71 | 15.72 / 2.69 | 15.70 / 2.72 | 15.75 / 2.86 | 15.78 / 2.85 |
| Replicates | 3 | 4 | 3 | 4 | 3 |
| Repeatability (avg. standard deviation of #D) | 0.2416 | 0.2724 | 0.1707 | 0.2696 | 0.2101 |

**Supplemental Table 2.**
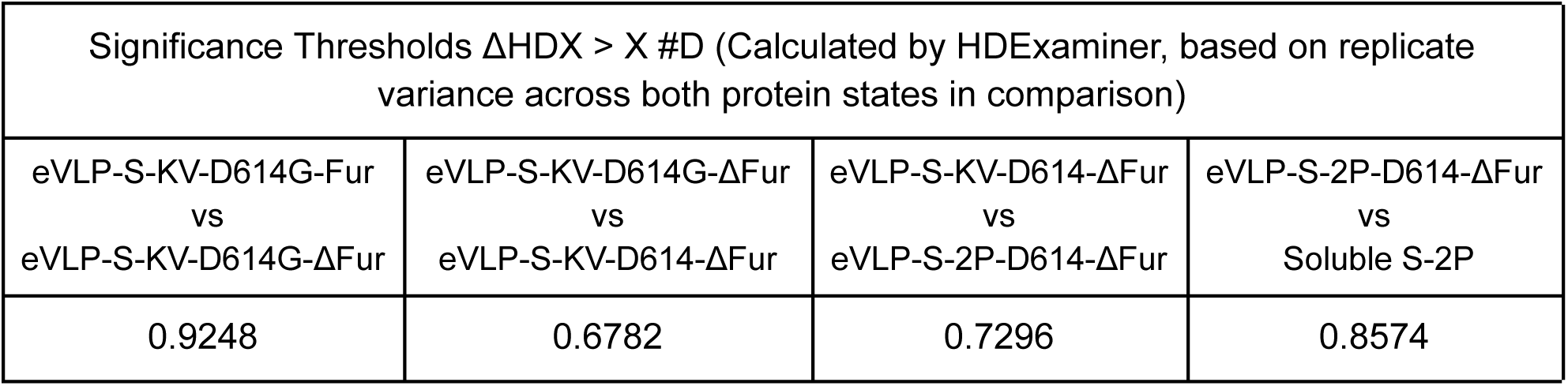

**Supplemental Figure 1.**
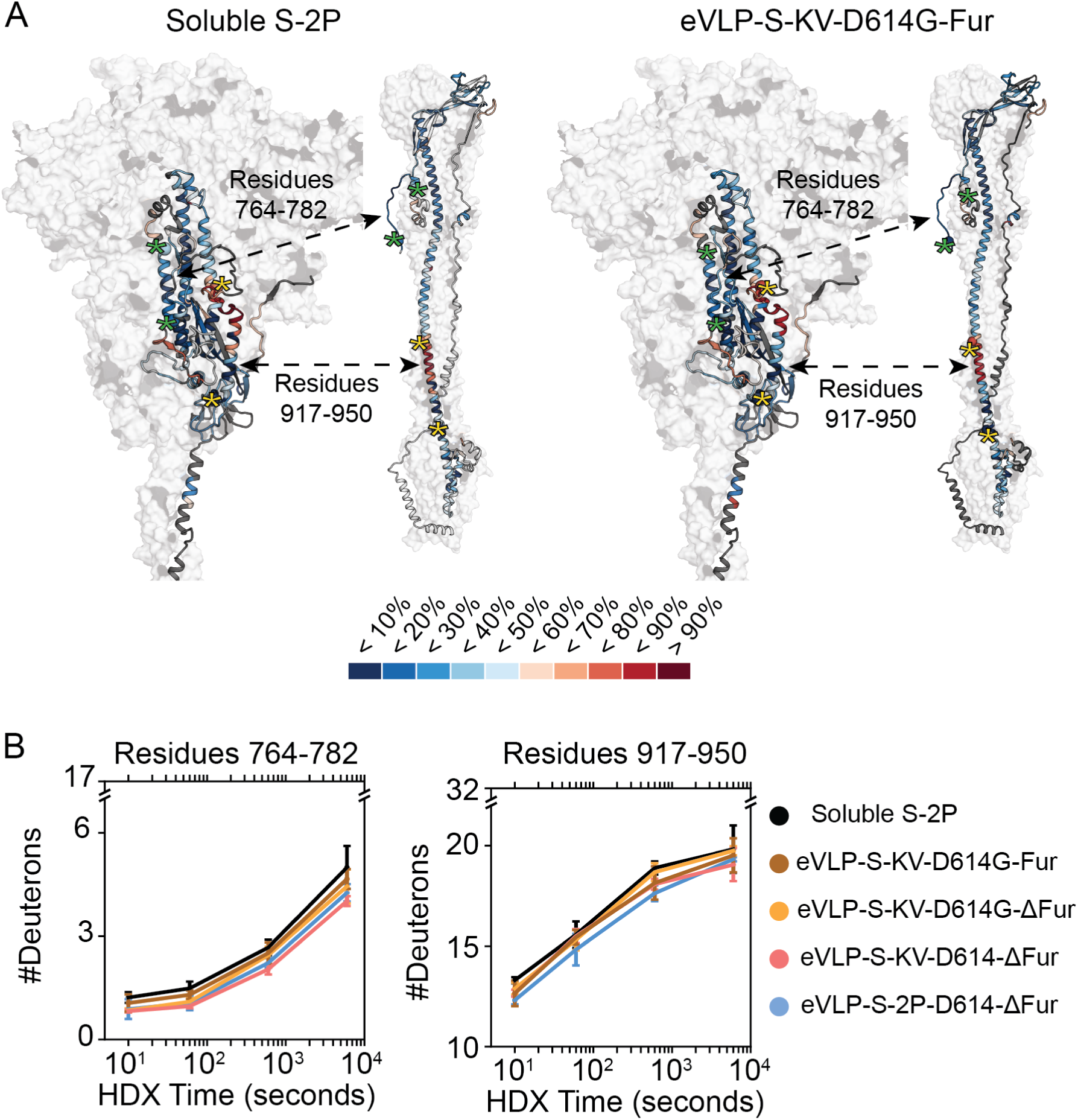
HDX-MS of all spike constructs is consistent with the prefusion conformation. (A) Heatmap of 60 seconds of exchange from S-2P (right) and eVLP-S-KV-D614G-Fur (left) mapped onto a single protomer of the S2 domain (in cartoon), the rest of the domains of the trimer are shown in a white surface on a model of the full-length prefusion structure ^28^ (left) and the postfusion structure (PDB ID: 8FDW) (right). Two regions that are in very different structural environments are denoted with asterisks. Residues 764-782 (green asterisks) which is in a core helix in the prefusion structure and an unstructured loop on the postfusion and residues 917-950 (yellow asterisks) which is solvent exposed and partially unstructured in the prefusion conformation while is helical and occluded from solvent in the postfusion. (B) Uptake plots from the two regions denoted by asterisks in part (A) for all constructs used in continuous labeling experiments. The exchange of these peptides is consistent with the prefusion structure and inconsistent with the post fusion structure. Furthermore while there are small differences in the magnitude of exchange, the overall shape of the uptake plot is the same indicating all the constructs have the same overall fold.

